# Fitness Effects of Somatic Mutations Accumulating during Vegetative Growth

**DOI:** 10.1101/392175

**Authors:** Mitchell B. Cruzan, Matthew A. Streisfeld, Jaime A. Schwoch

## Abstract

The unique life form of plants promotes the accumulation of somatic mutations that can be passed to offspring in the next generation, because the same meristem cells responsible for vegetative growth also generate gametes for sexual reproduction. However, little is known about the consequences of somatic mutation accumulation for offspring fitness. We evaluate the fitness effects of somatic mutations in *Mimulus guttatus* by comparing progeny from self-pollinations made within the same flower (autogamy) to progeny from self-pollinations made between stems on the same plant (geitonogamy). The effects of somatic mutations are evident from this comparison, as autogamy leads to homozygosity of a proportion of somatic mutations, but progeny from geitonogamy remain heterozygous for mutations unique to each stem. In two different experiments, we find consistent fitness effects of somatic mutations from individual stems. Surprisingly, several progeny groups from autogamous crosses displayed increases in fitness compared to progeny from geitonogamy crosses, indicating that beneficial somatic mutations were prevalent in some stems. These results support the hypothesis that somatic mutations accumulate during vegetative growth, but they are filtered by different forms of selection that occur throughout development, resulting in the culling of expressed deleterious mutations and the retention of beneficial mutations.

## Introduction

Mutation is the source of variation for evolution and adaptation, but organisms differ in whether mutations originating during gamete formation (meiosis) or somatic growth (mitosis) contribute to heritable variation. For the vast majority of organisms, including viruses, unicellular microbes, and some multicellular eukaryotes, sexual reproduction is rare or absent. In these organisms, mutations can occur during mitotic cell replication, and the primary mechanism for adaptation and diversification occurs via selection on cell lineages without recombination. By contrast, acquired mutations occurring during somatic growth of animals are not heritable. This is because in metazoans (but possibly excepting corals; Schweinsberg et al. 2014; Barfield et al. 2016), the germline is determined early in development and relatively few cell divisions occur before the formation of gametes (the Weismann Barrier; Buss 1983). Consequently, heritable mutations only occur in animals during the development of gonads and gametes.

Plants differ from animals and microbes, because mutations contributing to heritable variation can arise both during gamete formation and somatic growth (Antolin and Strobeck 1985; Klekowski and Godfrey 1989). This is due to the fact that plants lack a separate germline; the same germ cells within apical meristems subtend both vegetative growth and gamete development. Consequently, plants have the capacity to accumulate somatic mutations during vegetative growth that can be passed to the next generation (McKnight et al. 2002; Klekowski 2003; Schultz and Scofield 2009; Bobiwash et al. 2013; Dubrovina and Kiselev 2016; Watson et al. 2016; Schmid-Siegert et al. 2017; Yu et al. 2020). This aspect of plant biology is well known (Monro and Poore 2009; Ally et al. 2010; Reusch and Bostrom 2011), and somatic mutation accumulation has been important in agriculture, where the origin of many clonally-derived varieties of fruits, including citrus, apples, and wine grapes, have been cultivated by grafting from genetically differentiated bud tips (Aradhya et al. 2003; McKey et al. 2010; Miller and Gross 2011; Vezzulli et al. 2012; Jarni et al. 2015; Pelsy et al. 2015). However, there is disagreement over the extent and evolutionary importance of somatic mutation accumulation in plants (Schultz and Scofield 2009; Burian et al. 2016; Watson et al. 2016; Kuhlemeier 2017; Schmid-Siegert et al. 2017; Plomion et al. 2018; Hanlon et al. 2019).

In general, it is difficult to assess the effects of somatic mutations accumulating during growth because clonal cell lineages lack recombination, and the fitness effects of beneficial mutations can be muted by the co-occurrence of detrimental genetic variants (Hill-Robertson Effect; Selective Interference; Hill and Robertson 1966; Fogle et al. 2008). Although sexual reproduction does allow mutations to segregate among progeny, it is difficult to detect the fitness effects of somatic mutations in eukaryotes with separate sexes. This is because the genomes of two individuals are combined, so individual somatic mutations that occur in one individual will always be heterozygous in offspring (Long et al. 2015). A more effective approach would be to make self-fertilizations in a hermaphroditic species, so a proportion of offspring would be homozygous for novel somatic mutations. In this study, we make crosses within individual clones of hermaphroditic plants to assess the fitness effects of inherited mutations that accumulated during somatic growth.

Hermaphroditic plants have several advantages for the study of somatic mutations. Separate stems on the same plant contain multiple germ cell lineages that are derived from the same zygote, but they may differ for somatic mutations that have accumulated during stem elongation. To produce recombinant progeny segregating for somatic mutations unique to each stem, crosses can be made either within the same flower (autogamy), or between flowers on separate stems of the same plant (inter-ramet geitonogamy – hereafter referred to simply as geitonogamy; Fig. 1). These crosses are both self-fertilizations, but the offspring will differ in the complement of somatic mutations that they inherit. Mutations unique to a stem will be homozygous in 25% of autogamous progeny, while none of the progeny from geitonogamous crosses will be homozygous, and half will be heterozygous for unique somatic mutations. Thus, the average fitness effects of somatic mutations unique to each stem can be evaluated by comparing progeny generated by autogamous and geitonogamous crosses of the same stem. The potential for somatic mutations to accumulate during vegetative growth, and the ability to study their effects among recombinant progeny, make plants an attractive model for the study of fundamental processes of clonal evolution.

**Fig. 1.**
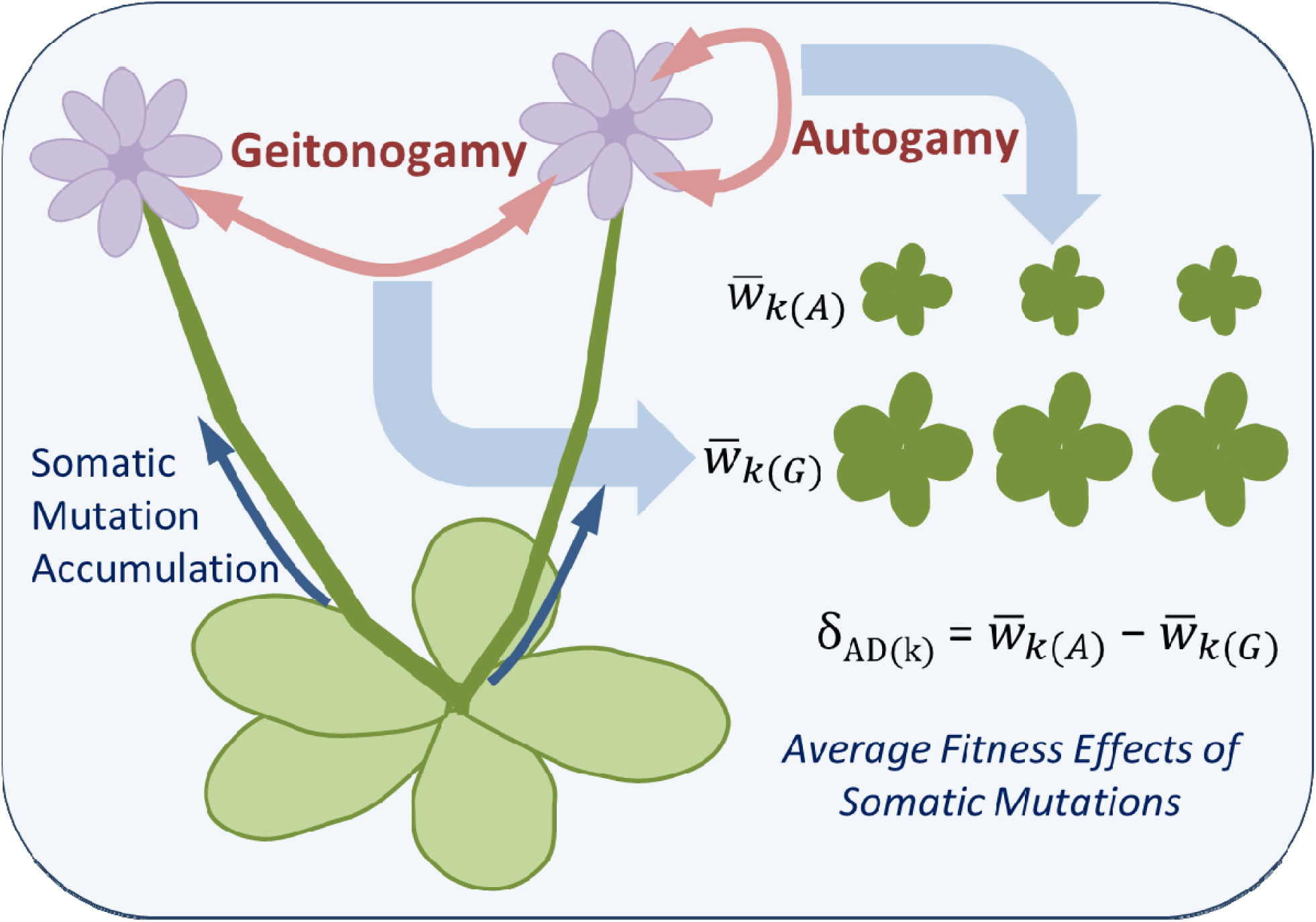
Experimental design to test for the average fitness effects of somatic mutations accumulating in stems during vegetative growth. See text for further explanation.

Plants grow from the division of a population of undifferentiated meristem cells within the stem tip that is known as the central zone. These germ cell lineages go on to produce future stem, leaf, and reproductive tissues (flowers). As plants grow, individual ramets of the same genet (separate stems or vegetatively propagated plants) can continue to become differentiated as they accumulate somatic mutations (Fig. 2). Therefore, when we consider the potential for meiotic and somatic mutations to contribute to the total mutational load of plant populations – particularly for long-lived plants – it becomes evident that not all of the mutations occurring during a plant’s lifespan are passed to the next generation (Cruzan 2018 pages 86-98). Indeed, plants have mechanisms of “developmental selection” (Buchholz 1922) that occur during vegetative growth and reproduction to filter the set of mutations that are inherited by progeny. For example, because new somatic mutations occur as a single copy within the diploid genome, their immediate fitness effects will depend on the combination of their fitness effects and their expression in the heterozygous state. Mathematical models have demonstrated that multiple unexpressed mutations (i.e. neutral and recessive deleterious mutations) are likely to accumulate as germ cells divide, so any selective cell deaths that occur during somatic growth will reduce the total mutational load and alter the composition of mutations carried by the germ cell population (Otto and Orive 1995; Otto and Hastings 1998). Thus, lineages that carry expressed deleterious mutations causing slower rates of division can be eliminated as they are replaced by cell lineages displaying more rapid growth (Otto and Orive 1995; Otto and Hastings 1998; Orive 2001; Elena and Lenski 2003; Greaves and Maley 2012; Long et al. 2015). (Fig. 2A). More rarely, expressed beneficial mutations will arise. Similar to the process of clonal evolution in mammalian cancer, any mutations that elevate the rate of cell division will tend to increase in frequency until they have replaced the entire germ cell population (clonal selective sweep; Nowell 1976; Lang et al. 2013; Fig. 2B green arrows). This process – referred to as cell lineage selection – will lead to some ramets containing a prevalence of deleterious somatic mutations, while others could possess beneficial mutations. The potential for these clonal selective sweeps to occur during vegetative growth has been confirmed by the observation of somatic mutations occurring at frequencies near 50% (heterozygosity) within the stem tissue (Yu et al. 2020; Schwoch et al. unpublished data). Hence, germ cell populations in plant apical meristems have the potential to undergo clonal evolution during vegetative growth.

**Fig. 2.**
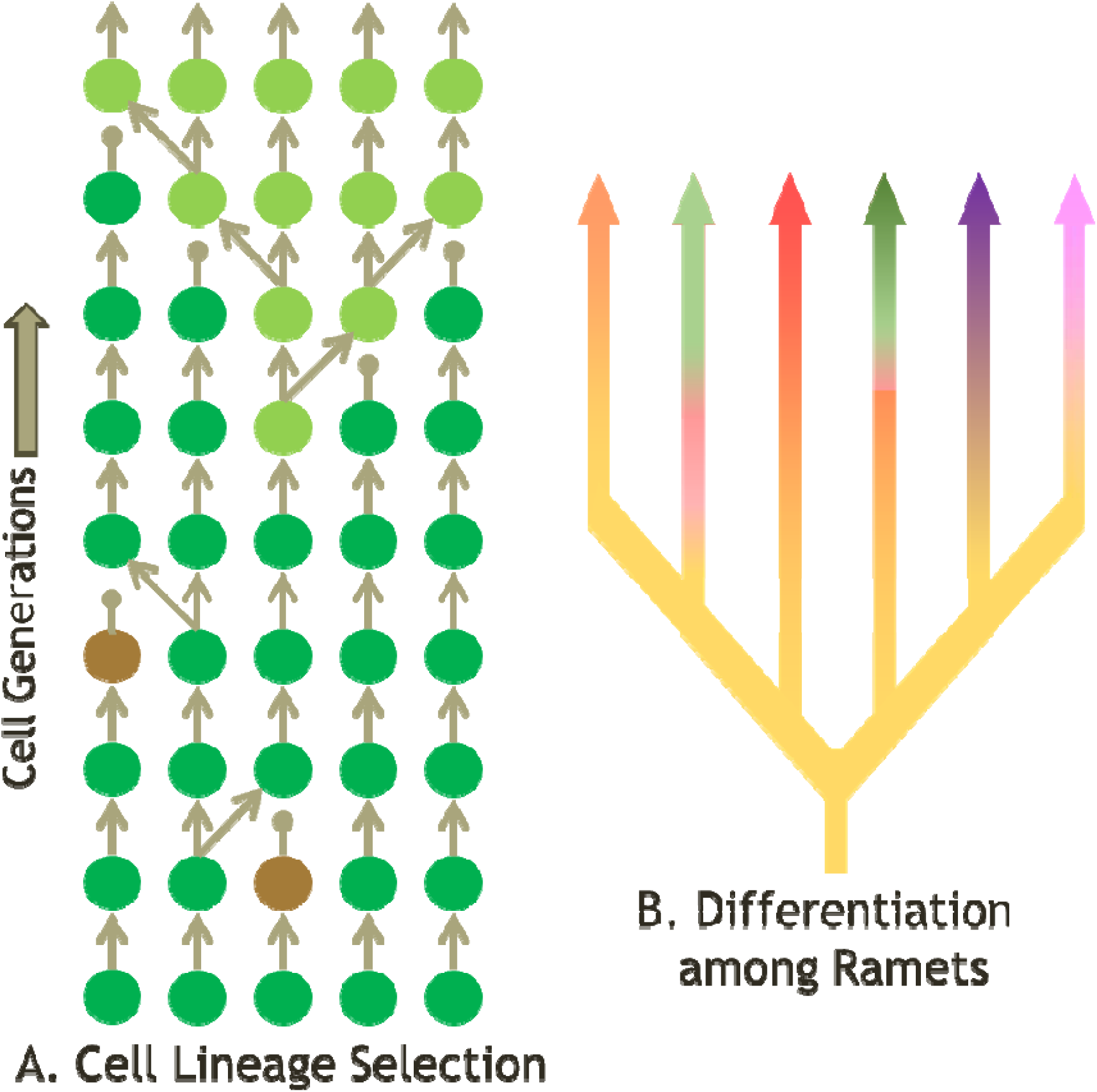
An illustration of cell lineage selection as it acts on a number of cell lines that differ in growth rate within the apical meristem of a plant. A) Expressed deleterious mutations are predicted to lead to the demise of a cell line (brown cells) that is replaced by neighboring cell lines. The appearance of a mutation that increases cell growth rate in the current environment results in a selective sweep of the meristem (light green cells). B) Multiple ramets of the same genet become differentiated by unique somatic mutations during vegetative growth. Most ramets carry deleterious mutations (orange, red, purple, and pink arrows), but expressed beneficial mutations cause clonal selective sweeps of the meristem in some ramets (green arrows).

In addition to cell lineage selection, some proportion of recessive deleterious mutations may be eliminated during the haploid life stage due to pollen tube attrition and pollen competition (Gemetophytic Selection; Mulcahy 1979; Cruzan 1989; Mable and Otto 1998; Armbruster and Rogers 2004; Arunkumar et al. 2013; Harder et al. 2016). Finally, a portion of deleterious mutations will be homozygous in zygotes, which can lead to higher rates of seed and fruit abortion (Selective Embryo Abortion; Husband and Schemske 1995; Korbecka et al. 2002), thereby increasing the average fitness of surviving offspring (Cruzan and Thomson 1997; Mena-AlÍ and Rocha 2005). The sum effects of these processes of developmental selection, which includes cell lineage selection, gametophytic selection, and selective embryo abortion, may filter the set of mutations that enters the next generation. Indeed, the effects of deleterious somatic mutations often are apparent as higher rates of embryo abortion after autogamous compared to geitonogamous pollinations, which is referred to as autogamy depression (Schultz and Scofield 2009). While autogamy depression for seed and fruit abortion has been observed in several species (reviewed in Bobiwash et al. 2013), no previous study has evaluated the fitness effects of somatic mutations inherited by progeny. The question we address here is whether somatic mutations that characterize individual ramets have measurable fitness effects in the offspring generation.

In this study, we use autogamous and geitonogamous self-pollinations to estimate the fitness effects of somatic mutations segregating in offspring of *Mimulus guttatus* DC (*Erythranthe guttata* G.L. Nesom; Phrymaceae). Populations of *M. guttatus* display a wide range of life histories – from annuals to herbaceous perennials that outcross to varying degrees (Wu et al. 2008). We use perennial *M. guttatus* plants that produce substantial vegetative growth prior to initiation of flowering. These plants are easily propagated from rosettes, and selfing produces substantial numbers of seeds to allow for statistical comparisons. In two separate experiments, we characterized the fitness effects of somatic mutations in perennial *M. guttatus* plants in controlled environments.

We begin by describing the analytical approaches we use, followed by providing specifics of the two experiments that we conducted. We show that somatic mutations accumulate during stem growth, as progeny from autogamous crosses have greater variance in mean fitness compared to progeny from geitonogamous crosses. We further reveal evidence for the presence of beneficial mutations in some stems, which indicates that developmental selection may be responsible for filtering mutations and modifying the distribution of fitness effects transmitted to progeny. The results from these experiments provide the first evidence that inherited somatic mutations can have fitness consequences for plants in the next generation, which suggests that somatic mutations may be an additional source of genetic variation that can be used for adaptation.

## Approach

We made autogamous and geitonogamous self-pollinations across multiple stems and estimated the fitness effects of somatic mutations segregating in progeny. To increase the chances of detecting somatic mutations that impacted fitness, we exposed plants to novel environments. According to Fisher’s geometric model, displacement from a fitness optimum increases the chances that any mutation would move a genotype in a direction that increases fitness (Fisher 1930; Stearns and Fenster 2016). Furthermore, we evaluated fitness of seedlings under novel selection regimes in the greenhouse. Exposing parental plants and seedlings to the same controlled environments is common in experimental designs that test for the fitness effects of mutations accumulating across generations (Shaw et al. 2002; Baer et al. 2007; Halligan and Keightley 2009), or in our case, for observing the fitness effects of somatic mutations arising within a single generation.

For a diploid plant, we can assume that somatic mutations (*a* → *a*’) will be in the heterozygous state when they first appear. For progeny arising by autogamy, a mutation will segregate as 25% homozygous (*a*’*a*’), 50% heterozygous (*aa*’), and 25% the original homozygote (*aa*). By contrast, because progeny from geitonogamous crosses will be segregating for mutations that are unique to each stem, half of them will be carrying mutations in the heterozygous state, and none of the progeny will be homozygous for mutations that arose in a single stem. The fitness effects of somatic mutations accumulating during vegetative growth have been quantified in terms of “autogamy depression” (*δ*_*AD*_) using the formula; *δ*_*AD*_ = 1 – *w*_*A*_*/w*_*G*_ (Schultz and Scofield 2009; Bobiwash et al. 2013), where *w*_*A*_ and *w*_*G*_ are the relative fitnesses of progeny derived from autogamous and geitonogamous crosses, respectively.

Tests of autogamy depression have been applied previously only to estimates of fitness based on fruit and seed set, and it generally assumes that fitness from autogamous crosses will be lower than from geitonogamy. However, somatic mutations that are transmitted to progeny can affect fitness in the offspring generation across different life stages, and beneficial mutations are possible. Thus, for each stem, we examined the fitness effects of somatic mutations unique to stem *k* as: 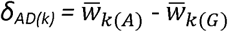, where 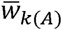 and 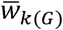 are the average fitnesses of the progeny from autogamous and geitonogamous crosses from stem *k*, respectively. We evaluate *δ*_*AD*_ for each stem separately, because we expect that individual stems will have unique complements of somatic mutations and thus differ in their fitness effects. By controlling for environmental effects with a common garden design, the parameter *δ*_*AD(k)*_ and its sign describe the average magnitude and overall direction of the fitness effects of all expressed somatic mutations that have been transmitted to the next generation. To evaluate the reliability of *δ*_*AD(k)*_ as an estimate of the fitness effects of somatic mutations, we simulated variation in the strength of selection and dominance of somatic mutations across multiple pairs of stems (ramets). We determined that values of *δ*_*AD(k)*_ were accurate across a range of values and assumptions for the effects of multiple mutations (Appendix 1).

To provide additional confirmation of the *δ*_*AD(k)*_ estimates, we developed an alternative approach for estimating the fitness effects of somatic mutations that was based entirely on the variance in mean fitness among autogamous progeny. Individual germ cell lineages within the meristem may possess unique sets of somatic mutations (Yu et al. 2020; Schwoch et al. in preparation), and flowers on the same stem can develop from only a portion of the germ cells present in the meristem (McManus and Veit 2002). Hence, no pair of flowers within a stem is likely to contain exactly the same complement of mutations. Therefore, we generated an additional estimate of the fitness effects of somatic mutations unique to a stem that is based on the variance in fitness of offspring from autogamous crosses. If somatic mutations affect offspring fitness, the variance in fitness should be greater for progeny groups from autogamy than from geitonogamy from the same stem, as long as mutations do not have complete expression in heterozygotes (i.e. *h* < 1.0; Appendix 2). Somatic mutations that are unique to stems will have different effects on the variance in fitness among progeny from autogamy depending on their fitness (*w*) and their expression in the heterozygous state (*h*). Our analytical results demonstrated that the large majority of variance in fitness among progeny from autogamy (as measured by the standard deviation, SD) could be explained by the fitness effects of a mutation rather than the dominance level (Appendix 2; Fig. S1). Moreover, the derived relationship allows us to estimate the fitness effects of mutations based on the standard deviation in fitness according to the equation: *w*_*SD*_ = *cSD*, where *c* is the slope of the linear relationship between *w*_*SD*_ and *SD*. For dominance levels of *h* = 0.0 and 1.0, *c* = 2.31, and the slope reaches a maximum of 3.01 when *h* = 0.5 (Appendix 2; Fig. S1). Thus, the variance in fitness among progeny from autogamous crosses provides a good approximation of the magnitude of the fitness effects (*w*) for somatic mutations accumulating in each stem. These two approaches are independent of each other, and therefore provide an important means to confirm our estimates of the average fitness effects of somatic mutations present in individual stems (Appendix 1 and 2).

From the considerations above, we can make three specific predictions that allow us to test for the fitness effects of somatic mutations that occur during stem growth and are passed to offspring. First, there should be greater variation among progeny groups from autogamy compared to those from geitonogamy (i.e. as indicated by the stem x cross type interaction for progeny fitness). Second, the average fitness of progeny from autogamy should be different from the progeny of geitonogamous crosses from the same stem. The difference in the fitness of progeny from autogamous vs. geitonogamous crosses provides an approximate estimate of the sign and magnitude of the average fitness effects of somatic mutations for individual stems. In addition to influencing the mean fitness, somatic mutations will increase the variance in fitness among progeny from autogamy. Consequently, the third prediction we can make is that the mean fitness effects of somatic mutations will scale linearly with the within-family standard deviation in fitness among progeny from autogamous crosses (Appendix 2). In general, we predict that if somatic mutations with fitness effects accumulate during vegetative growth, then the mean progeny fitness should differ between autogamous and geitonogamous crosses, and the variance in fitness of progeny from autogamous crosses would be greater than for progeny from geitonogamous crosses. Below, we test these predictions using two different experiments with *Mimulus guttatus*.

## First Experiment

### Methods

This experiment originally was designed to test the effects of somatic mutations on seed set and ovule abortion (autogamy depression; Schultz and Scofield 2009; Bobiwash et al. 2013), so we used multiple plants from several populations to obtain a large number of fruits. We grew plants of *M. guttatus* in the Research Greenhouse facility at Portland State University from seed collected in July 2013 from three different populations in northern Oregon (Jackson Bottom Wetlands–JB: 45.501794 N, - 122.98776 W; and two from Saddle Mountain–SMB: 45.9861 N, -123.6859 W, and SMC: 45.9634 N, - 123.6837 W). We assumed that greenhouse conditions were different enough from field environments to provide a novel selection regime. In August 2013, seeds were cold stratified on moist paper towels at 2°C for 30 days prior to being sown in soil. Seedlings were transplanted to pots (approximately 10 × 10 x 12 cm) and grown for seven months before the application of pollination treatments. Temperature was maintained between 21–26°C during the day, and 15–21°C at night. Supplemental HID lights ran for 12 hours a day when the seedlings first emerged, and 14 hours a day during adult growth.

After plants became established and began producing multiple stems, we conducted autogamous and geitonogamous self-pollinations using flowers on several stems (15 to 20 cm in length) from two plants from each of four maternal families representing each of the three populations. Flowers from pairs of stems on individual plants were reciprocally crossed (geitonogamy), or individual flowers from these same stems were self-pollinated (autogamy). A total of 139 pollinations were conducted across two treatments: limited (pollen was applied to stigmas with one touch from a plastic pipette tip) or excess (where the stigma surface was coated with pollen). Pollinations were conducted on 12 different days (pollination date) over several weeks in July 2014. Mature fruits were collected and placed individually into paper envelopes, and their contents were examined under a Leica MZ-16 stereoscope. The first 100 ovules from each fruit were categorized as filled seeds (brown, almond-shaped), unfertilized ovules (small, flattened and light-colored), or aborted (larger than unfertilized, dark-colored, shriveled). Ovules that were flattened and appeared to lack endosperm were assumed to be unfertilized or aborted and were not used in germination tests. Seed set and ovule abortion were analyzed using ANOVA models with the GLM procedure of SAS (SAS 2008), with population, maternal plant nested within population, and pollination date as random effects, and cross type (autogamous or geitonogamous) and pollination treatment as fixed effects. Seed set and ovule abortion data were approximately normal so were not transformed prior to analysis.

We assessed the fitness of progeny arising by autogamy and geitonogamy in the same greenhouse environment that was used to grow the parental plants. Seedlings from a subset of ten maternal plants that had fruits from both cross types and at least 20 filled seeds were sown in soil and transplanted to 36-cell trays (blocks) in September in a randomized incomplete block design. After three months of growth, the progeny were scored for survival, and above ground biomass was measured after drying at 60°C for at least 24 hours. Since all seedlings germinated within a few days of each other, the final biomass is an estimate of growth rate. The fitness of each progeny was estimated as its final biomass, which was log transformed and weighted by the survival frequency of progeny from the same cross. Growth rate is considered to be an appropriate estimate of fitness for perennials (Younginger et al. 2017). Furthermore, we evaluated fitness of seedlings under novel selection regimes in the greenhouse rather than under field conditions, which allowed us to control for environmental differences by having parental plants and seedlings exposed to the same conditions. These estimates were rescaled relative to the maximum value from all crosses, so that 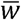 ranged from 0 to 1 across progeny from all crosses. Data were analyzed to estimate the mean fitness for progeny from autogamy 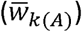 and geitonogamy 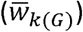 from each stem, and the variance in fitness (as measured by the standard deviation; SD) for each group of progeny produced by autogamy and geitonogamy from single stems.

### Results

Levels of seed set and germination were similar between cross types, but ovule abortion was greater in autogamous crosses. The mean number of seeds produced after autogamous (56.72 ±2.24, N = 87) and geitonogamous (57.52 ±3.42, N = 52) pollinations was similar (Cross in Table S2), and there was little variation among populations (Pop in Table S2), maternal plants nested within population (Mat(Pop) in Table S2), and pollination dates (Date in Table S2). Limited pollinations produced fewer seeds than excess pollinations (Pollen in Table S2), but the effect of pollen dosage in the overall model was not significant. However, the mean number of aborted ovules was greater for autogamous (17.36 ±1.27, N = 87) than for geitonogamous (13.90 ±1.70, N = 52) pollinations (Cross in Table S3), which is consistent with the predicted effects of autogamy depression (Schultz and Scofield 2009; Bobiwash et al. 2013). There also were significant differences among populations in the level of ovule abortion (Pop in Table S3), which perhaps reflects differences in historical inbreeding and levels of genetic load. There were significant differences in ovule abortion among maternal plants nested within population (Mat (Pop) in Table S3), but not for pollination treatments (Pollen in Table S3) or pollination dates (Date in Table S3). The overall model for ovule abortion was significant (Table S3). Seed germination was similar between autogamous (mean = 13.03 out of 20 planted per fruit) and geitonogamous crosses (mean = 13.28; F_1,101_ = 0.16, P = 0.694). Of the 354 seedlings that germinated, 202 survived, and interestingly, survival was higher for progeny from autogamy (67%) compared to geitonogamy (50%; chi-square = 12.03, P = 0.0005). For the surviving plants that flowered by the end of the experiment, flower production was correlated with growth rate (*r* = 0.58, P < 0.001, N = 282).

The average fitness of progeny from autogamy and geitonogamy varied widely among individual stems. Consistent with our first prediction for the effects of somatic mutations on offspring fitness, there was a significant interaction between cross type and stem identity (Table S4). Little of the variation in progeny fitness was explained by differences among stems (Mat in Table S4), but larger amounts were explained by cross type (Cross in Table S4) and among blocks (Tray in Table S4). When data were analyzed separately for each cross type, we found that mean fitness of progeny from autogamy varied among stems to a much greater degree (F_8,130_ = 4.44, P < 0.0001) than progeny from geitonogamous crosses (F_8,70_ = 1.47, P = 0.1866); a result that supports our first prediction for the effects of somatic mutations on progeny fitness. These data are analyzed further along with data from the second experiment described below.

## Second Experiment

The observation of greater variation in fitness among stems for progeny from autogamous crosses in the first experiment is consistent with the hypothesis that somatic mutations arising during vegetative growth are transmitted and can have substantial fitness effects in the next generation. Furthermore, the results from the first experiment do not support the view that somatic mutations during vegetative growth are rare (Burian et al. 2016; Groot and Laux 2016; Watson et al. 2016; Kuhlemeier 2017; Schmid-Siegert et al. 2017), but they are consistent with several studies demonstrating that somatic mutations can be inherited by offspring in the next generation (Klekowski and Godfrey 1989; Schultz and Scofield 2009; Bobiwash et al. 2013). However, the first experiment was not designed specifically to compare the effects of autogamous and geitonogamous pollinations on offspring fitness. Thus, to confirm results from the first experiment, we performed a second experiment to provide a more robust assessment of the effects of autogamous and geitonogamous self-pollination on the fitness of offspring. The second experiment used a single clone of *M. guttatus* to make comparisons between autogamous and geitonogamous pollinations paired at the same node. The plant chosen was a self-compatible perennial that displayed vigorous vegetative growth. We exposed plants to novel conditions using high salinity under hydroponics cultivation, which increased the potential that mutations would have fitness consequences. Moreover, our recent genomic results confirm more low-frequency somatic mutation among plants exposed to high salt stress compared to the control hydroponics treatment without added salt (Schwoch et al. in preparation). This experimental design provided a more refined test of the hypothesis that the fitness effects of somatic mutations differed in the offspring of autogamous and geitonogamous crosses.

### Methods

To assess the fitness effects of mutations that accumulated during vegetative growth, a single plant (genet BV, obtained from Willamette Gardens native plant nursery, Corvallis, OR) was vegetatively propagated to generate 12 plants (ramets) that were exposed to high salinity and control conditions. Plants were grown in pea gravel (4–8 mm) in pots placed in four 53 L tubs using a flood and drain hydroponics system (flooding at 15 min intervals). Two tubs had no added salt and were used as controls, and two tubs had high salinity. The initial salt concentration in the high salinity treatment was 5 mM but increased weekly to 25 mM after plants became established. Salt concentrations were monitored using a conductivity meter to ensure stable concentrations. To provide nutrients, 30 ml of hydroponics fertilizer (FloraGrow, Planet Natural, Bozeman, MT) was added per tub. During the course of the experiment, some plants grew substantially. We imposed selection to favor the fastest growing ramets over the next three months by repeatedly removing single rosettes and transplanting them back into the hydroponics system up to four times.

To promote stress recovery, plants were transplanted to soil for six months, which included a two-month vernalization period in a growth chamber (4°C and 8 h light; Conviron E8, Controlled Environments Ltd., Winnipeg, Manitoba, Canada). After vernalization, plants were returned to the greenhouse to induce flowering. Autogamous and geitonogamous pollinations were made to pairs of flowers at single nodes or consecutive nodes (seven nodes and 14 pollinations total) on the largest ramets in each of the control and high salt treatments. To account for somatic mutation turnover that may occur due to the effects of cell lineage selection during stem growth, we compared progeny from autogamous and geitonogamous pollinations at pairs of flowers from the same node. Without a priori knowledge of the expression of somatic mutations in heterozygotes, it is difficult to determine the best pollen donor for geitonogamous crosses. Consequently, we opted to generate progeny from geitonogamy by pollinating flowers with pollen from a potentially more genetically divergent ramet from the other treatment (i.e. salt pollinated with control pollen, and control pollinated with salt pollen). The fruits were collected, and the total number of aborted and mature seeds and unfertilized ovules were counted under a dissecting microscope. Seeds were planted in soil in trays with three seeds per cell. Seeds were cold stratified in moist soil for three weeks before they germinated in the greenhouse.

To determine whether seedlings from autogamous crosses from ramets exposed to salt stress showed improved performance under the same conditions, all progeny were exposed to high salinity. After germination and establishment in soil, seedlings from autogamous and geitonogamous crosses from control and salt stress ramets were transplanted into pots filled with pea gravel and subjected to high salt in the hydroponics system, as described above. A total of 239 seedlings from 11 fruits (five autogamous and six geitonogamous) were randomly and evenly distributed among 12 hydroponic tubs to ensure equal representation across blocks (tubs). Plant size was measured as the product of the length and width of vegetative spread after two months of growth and was used as a proxy for overall plant performance. Since plants germinated within a few days of each other, plant size represents a good estimate of growth rate. Salt concentration increased from 10 mM - 37.5 mM over the course of the experiment to induce mortality (∼57% across all progeny groups). Fitness was estimated as plant size (log transformed to improve normality) weighted by the survival frequency for progeny from the same cross.

### Results

Similar to experiment 1 and consistent with our first prediction for the effects of somatic mutations, there was greater variation in the fitness of progeny from autogamous compared to geitonogamous crosses, as indicated by a significant interaction between cross type and stem/node identity (Stem/Node*Cross in Table S5). There was no consistent difference in progeny fitness between autogamous and geitonogamous pollinations when averaged across stems (Cross in Table S5), but there were larger differences among individual stems/nodes (Stem/Node in Table S5). Also consistent with our first prediction, there was significant variation in progeny fitness among stems (F_2,89_ = 3.76, P = 0.028) and nodes (nested within stems; F_3,89_ = 2.59, P = 0.0596) for autogamous, but not geitonogamous crosses (F_1,136_ = 0.63, P = 0.431 and F_4,136_ = 1.19, P = 0.317 for stems and nodes, respectively). There were significant differences in average size among the blocks (Tray in Table S5), but there was no consistent difference in the performance of progeny based on the treatment history of their maternal ramets (F_1,126_ = 0.02, P = 0.888), and there was no interaction between cross type and historical treatment (F_1,136_ = 0.64, P = 0.423). Three nodes were dropped from further analysis, because one of the paired fruits aborted or produced fewer than five seedlings. However, similar to experiment 1, and consistent with the hypothesis that somatic mutations accumulate during stem growth, the variance in fitness among progeny from autogamy among stems was greater than for geitonogamy. Since results from the two experiments were similar, the data for four nodes from experiment 2 were analyzed together with the first experiment, as described in the next section.

## Fitness Estimates

Results from the two experiments described above are consistent with the hypothesis that somatic mutations unique to individual stems can have demonstrable effects on the fitness of progeny when a proportion is made homozygous by autogamous self-pollination. In both experiments, the variance in fitness among progeny groups from autogamy was significantly greater than among progeny from geitonogamy from the same set of stems. As the results from these two experiments are qualitatively similar, we estimated the fitness effects of somatic mutations in each stem (ten comparisons of autogamy and geitonogamy from Experiment 1 and four from Experiment 2, for a total of 14 comparisons). Specifically, we tested the second and third predictions described above. Since 25% of somatic mutations unique to a stem will be homozygous in autogamous progeny, we expect that their fitness effects will be greater than for progeny from geitonogamous crosses on the same stem (providing an estimate of *δ*_*AD(k)*_). In addition, the within-family variance in fitness among progeny should be greater after autogamy compared to geitonogamy (providing an estimate of *w*_*SD*_). These estimates of the fitness effects of somatic mutations should be similar, so we assessed whether there was a positive relationship between the variance in fitness among progeny from single fruits produced by autogamous pollination (*w*_*SD*_) and the fitness effects of somatic mutations unique to each stem estimated from the average fitness of progeny from autogamous and geitonogamous crosses to the same stem (*δ*_*AD(k)*_).

Estimates of the average fitness effects of somatic mutations unique to each stem were made based on the fitness of progeny from autogamous 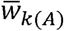 and geitonogamous 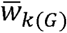 crosses (Table S1 in Appendix 3). In both experiments, fitness was estimated as growth rate weighted by the survival of seedlings from the same fruit and scaled to a maximum value of 1.0. We have demonstrated that growth rate is correlated with flower production, and it has been shown to be a good predictor of fitness among perennial plants (Younginger et al. 2017). The average fitness effects of mutations (*δ*_*AD(k)*_) for each stem (first experiment) or stem/node combination (second experiment) were calculated as the difference in fitness between progeny from autogamous and geitonogamous crosses, as described above 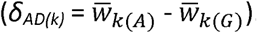. We observed extensive variation in *δ*_*AD(k)*_ among stems, with nine stems that were significantly different from zero (Fig. 3). Four of the stems had average fitness effects of somatic mutations that were positive, suggesting a net beneficial effect of somatic mutations transmitted to offspring. In addition, the average fitness of progeny from autogamous and geitonogamous crosses on the same stems or nodes was positively correlated (Fig. S2; Table S6). Note also that the *δ*_*AD(k)*_ estimates that deviated the most from zero generally had the highest (or lowest) 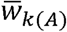 values relative to all of the stems (Fig. S3). However, this was not always the case, as one stem (stem 9) with a value of *δ*_*AD(k)*_ close to zero produced progeny with relatively high fitness after both autogamous and geitonogamous pollination.

**Fig. 3.**
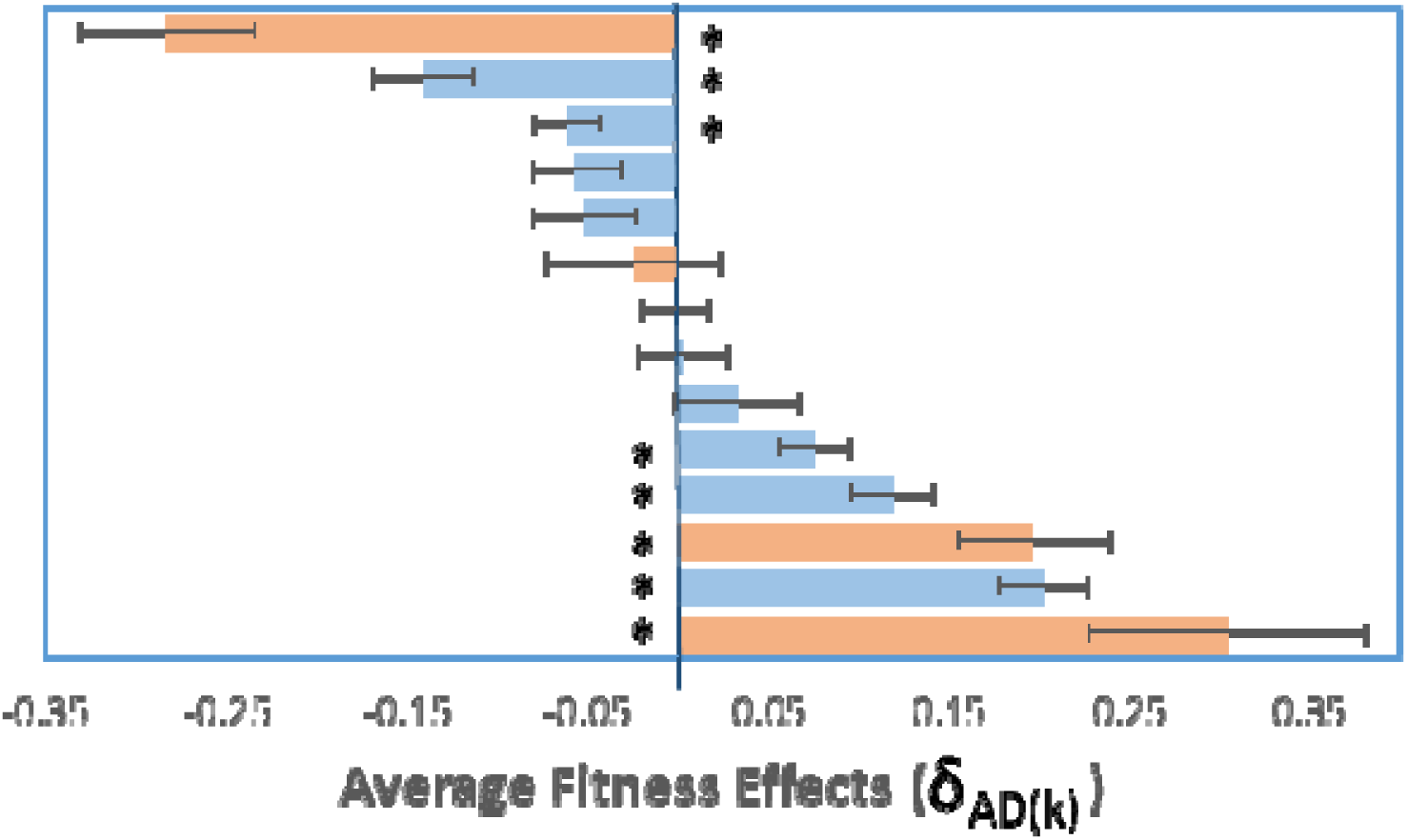
Estimates of *δ*_*AD(k)*_ for fourteen different stems (ramets) from two separate experiments (Experiment 1–blue bars; Experiment 2–orange bars) based on mean progeny fitness after autogamous 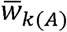 and geitonogamous 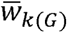 self-pollinations 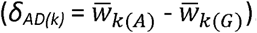. Horizontal lines represent standard errors. Asterisks indicate values of *δ*_*AD(k)*_ that are significantly different from zero based on the t value, calculated as 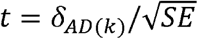 with *n*-1 df, where n is the mean of sample sizes for progeny from autogamy and geitonogamy. The relationship between 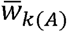 and 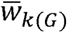 across stems is shown in Fig. S2. Means and sample sizes for progeny groups are available in Table S6 and Fig. S3.

To make an independent estimate of the average fitness effects for mutations unique to each stem or node, we used the standard deviation in progeny fitness from autogamous crosses based on the relationship *w*_*SD*_ *= cSD*, where *c* = 3, which assumes dominance close to *h* = 0.5 (qualitatively similar results are obtained if we choose other values of c between 3 and 2.31; Appendix 2). Because the sign of *w*_*SD*_ could not be inferred directly from this approach, we used the sign estimated from the *δ*_*AD(k)*_ method (Fig. 4; Appendix 2). Our simulations show that the sign of *δ*_*AD(k)*_ is almost always estimated correctly (Appendix 1). For stems with values of *δ*_*AD(k)*_ greater than zero, there was a strong positive relationship between *δ*_*AD(k)*_ and estimates of *w*_*SD*_ made from the within-family variation among progeny from autogamy (Fig. 4). In contrast, the relationship for negative values of *δ*_*AD(k)*_ appeared to be driven largely by a single observation. Note that this observation was not supported by a similarly high value of *w*_*SD*_, and the remaining negative fitness effects were more modest based on both estimates. This one highly negative estimate of *δ*_*AD(k)*_, may be due to a large influence of mutations from the second stem on the fitness of geitonogamous progeny (note that this stem had the highest estimate of 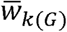; Fig. S2). It is also notable that the variance in fitness for progeny from autogamy did not decline to zero for values of *δ*_*AD(k)*_ close to zero, which could be due to the presence of both beneficial and detrimental mutations, and possibly genetic background effects (i.e. epistasis), but it may also reflect uncontrolled environmental variation. Overall, the results from both approaches reveal consistent estimates of the average fitness consequences associated with the accumulation of somatic mutations in individual stems.

**Fig. 4.**
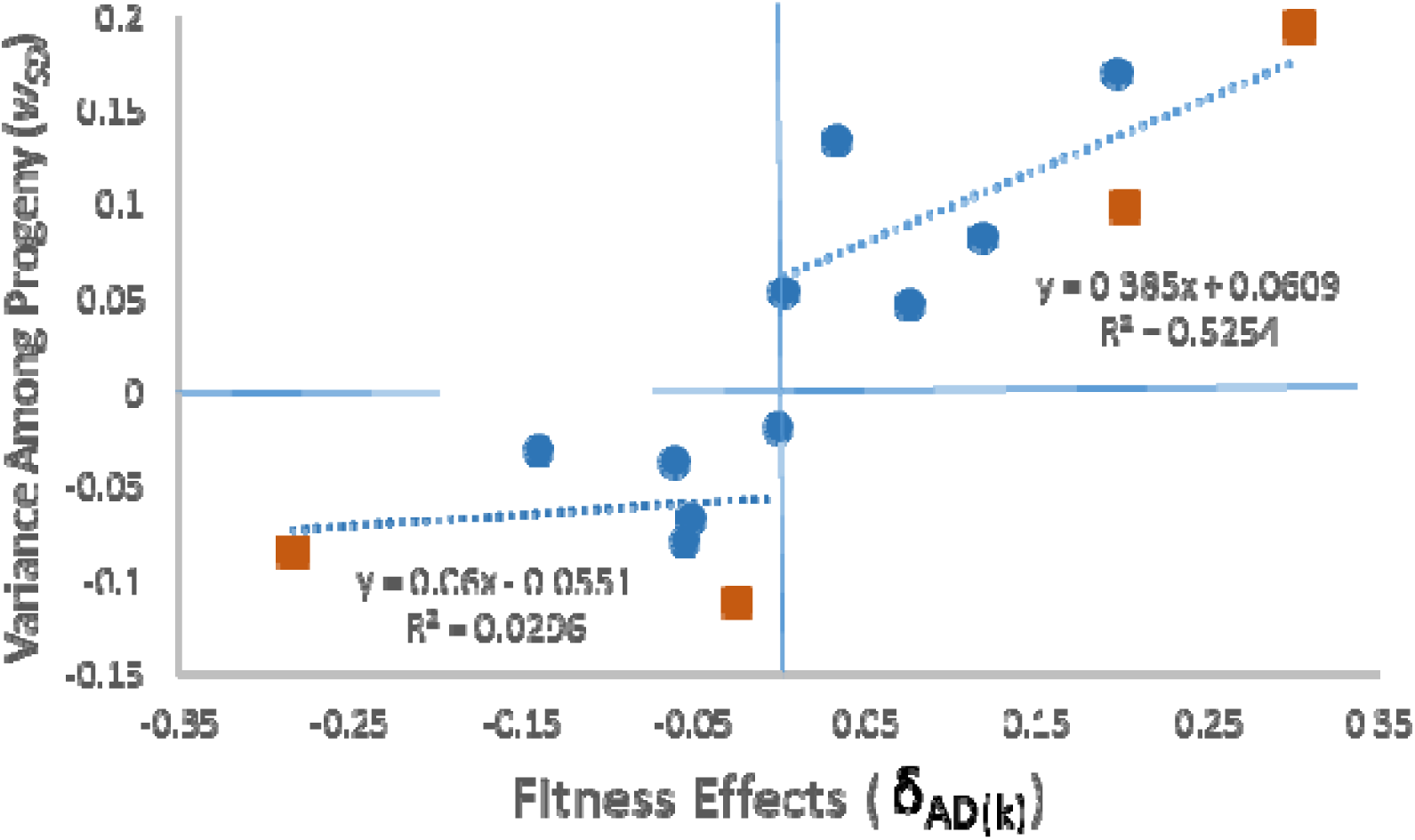
Relationships between estimates of fitness effects of somatic mutations, based either on the difference in fitness of progeny from autogamy and geitonogamy (*δ*_*AD(k)*_), or the standard deviation in fitness within progeny groups from autogamy for each stem (*w*_*SD*_). Estimates of *w*_*SD*_ corresponding to negative values of *δ*_*AD(k)*_ were transformed to negative values. Estimates from Experiment 1 are indicated by blue circles and from Experiment 2 are orange squares. Dashed lines indicate the separate relationships for positive and negative values of fitness estimates.

## Discussion

In this study, we have shown that somatic mutations can accumulate during vegetative growth and have substantial fitness effects on plants in the next generation. Consistent with previous work (Bobiwash et al. 2013), we observed significant autogamy depression in the form of greater ovule abortion in autogamous relative to geitonogamous crosses. However, we also detected more variation in survival and growth rate for progeny from autogamous compared to geitonogamous self-pollinations, which provides the first evidence that somatic mutations that accumulated during vegetative growth can have demonstrable effects on the fitness of plants in the next generation. Both the differences in the mean fitness between progeny from autogamy and geitonogamy (*δ*_*AD(k)*_) and variation in fitness of autogamous progeny (*w*_*SD*_) provided consistent estimates of the average fitness effects of somatic mutations segregating in progeny. We found evidence for the effects of beneficial mutations in progeny from some stems, with estimates of *δ*_*AD(k)*_ and *w*_*SD*_ exceeding 0.1 in four cases, while estimates for negative fitness effects were more modest (mostly > -0.15). Furthermore, the observation that progeny from autogamous crosses occasionally had higher fitness than progeny from geitonogamous crosses argues for the accumulation of beneficial mutations and implies that many deleterious mutations are culled by the various types of developmental selection prior to seed dispersal. These results imply that even though numerous somatic mutations are generated during vegetative growth, many of them are filtered due to different forms of developmental selection, which results in a shift in the distribution of fitness effects for the mutations that are passed to the next generation.

Somatic mutations accumulating during vegetative growth had an overall positive effect for four of the stems tested. This result is contrary to widely held views that the accumulation of beneficial somatic mutations should be exceedingly rare (Crow 1993; Charlesworth and Willis 2009). However, an explanation for these findings is afforded by the unique biology of plants; somatic mutations in germ cells in the central zone of the apical meristem could contribute to higher rates of division for some cell lineages over others (Otto and Orive 1995; Otto and Hastings 1998). This hypothesis is supported by the observation that a minority of somatic mutations occur at high frequencies in meristem tissue (Yu et al. 2020; Schwoch et al. unpublished data), suggesting that clonal selective sweeps have occurred during stem elongation. Thus, cell lineage selection has the potential to retain beneficial somatic mutations, as individual cell lineages divide at faster rates resulting in clonal evolution during vegetative growth.

Although there may be few opportunities for beneficial changes to alter basic cellular metabolism, it is becoming apparent from experimental evolution studies with microbes that even basic aspects of cellular metabolism can be sensitive to environmental conditions, which can increase the chances that mutations in clonal populations are beneficial (e.g., Lang et al. 2013; Lee and Marx 2013; Maharjan et al. 2015). In this regard, cell lineage selection in a plant meristem represents a potentially powerful forum for the removal of deleterious somatic mutations while favoring the retention of beneficial ones. In addition, gametophytic selection and selective embryo abortion can act as prominent additional filters (Mulcahy 1979; Cruzan 1989; Mable and Otto 1998; Armbruster and Rogers 2004; Arunkumar et al. 2013; Harder et al. 2016), but they are most likely to have effects by culling deleterious mutations. Regardless, the combined effects of these different forms of developmental selection appear to have had a considerable effect on filtering of somatic mutations under controlled conditions in the greenhouse, such that the distribution of fitness effects among stems has shifted to include more beneficial mutations than expected. Future studies should test whether similar findings are found under field conditions, which could indicate a prominent role for somatic mutation in local adaptation.

An alternative explanation for the observed fitness differences is that exposure to environmental stress has induced heritable epigenetic modifications (Quadrana and Colot 2016). However, epigenetic modifications are considered to be consistent in direction and predictable, as they are hypothesized to represent an adaptive response to historic exposure to similar stressors (Baulcombe and Dean 2014; Itabashi et al. 2018). The results of the experiments described here are unlikely to support a role for epigenetics, because fitness responses in the next generation were inconsistent in direction and magnitude, and they were not predictable based on environmental exposure of the parent stem. Indeed, among stems and nodes, the mean fitness of progeny from autogamy displayed both increases and decreases compared to the geitonogamy controls, which is consistent with the hypothesis that individual ramets are accumulating unique complements of somatic mutations. Furthermore, these conclusions are supported by the observation of numerous somatic variants in the transcriptomes of meristem tissue from multiple ramets derived from a single genet (Yu et al. 2020; Schwoch et al. unpublished data). Thus, the results of the current study appear to indicate that somatic mutations accumulating during stem growth are responsible for the observed fitness effects.

The potential for the acquisition of mutations during vegetative growth is a well-known aspect of plant biology (Klekowski 2003; Schultz and Scofield 2009; Bobiwash et al. 2013), but no previous study has demonstrated the effects of somatic mutations on the fitness of progeny in the next generation. Moreover, most studies have focused on the detrimental effects of somatic mutations; chloroplast mutants have been observed in a number of species (Klekowski 2003), declines in pollen fertility were found in older clones of quaking aspen (Ally et al. 2010), and higher rates of seed and fruit abortion after autogamous pollinations were found in several studies (reviewed in Bobiwash et al. 2013). In contrast, agriculturalists have taken advantage of beneficial somatic mutations to improve economically important plants (Aradhya et al. 2003; McKey et al. 2010; Miller and Gross 2011; Vezzulli et al. 2012; Jarni et al. 2015; Pelsy et al. 2015), and a handful of studies report phenotypic responses to selection in asexual lineages. For example, Breese et al. (1965) succeeded in selecting for increased tillering ability (production of new grass stems) within genets of perennial ryegrass (Lolium perenne). Similarly, artificial selection on clonal lineages effectively improved branching in the red seaweed, Asparagopsis armata (Monro and Poore 2009). The current study on *M. guttatus* contributes to this literature by highlighting the potential for plants to exhibit significant levels of clonal variation within a single generation.

The transmission of beneficial mutations in autogamous crosses may explain some heretofore difficult to understand results from mutation accumulation studies. Our results suggest that beneficial somatic mutations are likely to be partially dominant (i.e. not completely recessive), because they would have to be expressed in the heterozygous state to be favored by cell lineage selection. Mutations accumulating during vegetative growth could contribute to standing genetic variation, but autogamy may be more effective for the accumulation of beneficial somatic mutations in populations than geitonogamy or outcrossing, because beneficial mutations could be made homozygous in progeny in a single generation. Depending on the crossing design, outcrossing would take at least two generations for beneficial somatic mutations to be made homozygous, and under geitonogamy they would remain heterozygous. It is striking that high rates of beneficial mutation accumulation have been observed in at least some mutation accumulation studies in the autogamous plant Arabidopsis thaliana (Shaw et al. 2002; Rutter et al. 2010; Rutter et al. 2012), but not in outcrossing and partially-selfing species of Amsinckia (Schoen 2005). With a few exceptions (e.g., Baer et al. 2005; Denver et al. 2010), nearly all mutation accumulation studies on animals consistently show a prevalence of deleterious mutations (Baer et al. 2007; Halligan and Keightley 2009). Similarly, our results suggest that the adaptive potential of autogamous plants may be greater than previously thought, which may help explain the wider geographic ranges of selfing compared to closely-related outcrossing species (Grossenbacher et al. 2015; Grant and Kalisz 2020). Although developmental selection has the potential to contribute to adaptation in all plants, its effects may be enhanced in autogamous lineages, because beneficial mutations arising during vegetative growth have a greater chance of becoming homozygous in offspring and being retained across generations.

As stems grow, mutations can be generated during every mitotic cell division, so the potential for somatic mutation accumulation in plants appears substantial. Thus, understanding how long-lived plants, such as trees, avoid mutational meltdown from the accumulation of deleterious somatic mutations remains a longstanding question. Paradoxically, however, the rate of mutation accumulation observed across generations in plant and animal genomes is similar (Gaut et al. 2011). One hypothesis for this pattern is that, similar to animal germlines, a subset of cell lines in the apical meristem undergoes fewer mitotic divisions, which would protect lineages from the negative effects of mutation accumulation during development of the soma (Plant Germline Hypothesis; Burian et al. 2016; Cruzan 2018; page 90). An alternative hypothesis posits that somatic mutations are generated in apical meristems during plant growth, but these mutations are filtered by developmental selection prior to the establishment of offspring (Somatic Mutation Accumulation Hypothesis). Because plants have retained the capacity to undergo clonal evolution from their algal ancestors, the ability to filter mutations during growth and reproduction has existed for some time, and thus developmental selection has the potential to skew the distribution of fitness effects of transmitted mutations to include a larger proportion of beneficial mutations than would be expected through random processes. In addition, this provides a reasonable explanation for why longer-lived plants appear to have slower rates of mutation accumulation across generations (Yue et al. 2010; Gaut et al. 2011). This could be due to the fact that longer generation time leads to more time between recombination events, which can lead to more background selection in non-recombining cell lineages during vegetative growth (Cruzan 2018; pages 94-95). Recent studies in oaks, spruce, and eel grass (Plomion et al. 2018; Hanlon et al. 2019; Yu et al. 2020), as well as our unpublished data from *M. guttatus*, confirm that multiple somatic variants are likely, even in plants of very different size. Moreover, results from the current study identify the presence of beneficial mutations that were passed to offspring in the next generation, likely a consequence of developmental selection. Future work that combines information from experiments evaluating the genomic consequences of somatic variation with anatomical estimates of stem cell population dynamics will allow for the development of new models that provide insights into the extent and limitations of somatic evolution in plants.

In conclusion, despite the potential for somatic mutation accumulation to generate novel genetic variation in plant populations, its role in evolution remains almost entirely unexplored. Our estimates of the fitness effects of somatic mutations were consistent across two different methods and indicate that some stems possessed a prevalence of deleterious mutations while others produced autogamous progeny with high fitness, which indicates the presence of beneficial mutations. Even though high levels of mutation accumulation are often believed to be detrimental, the basic biology of plants suggests that the role of somatic mutations in plant evolution should be considered carefully in the future. Moving forward, our results indicate that we must keep in mind that all eukaryotes are not necessarily equivalent, and specific features of the organism’s biology may have unexpected consequences for the evolutionary phenomena that we observe. Future lines of investigation will improve our understanding of these fundamental aspects of plant development and evolution that may have contributed to the remarkable diversification of plants, and may help to account for some of the extensive variation in mutation rates detected among lineages.

## Acknowledgements

We thank J. Thompson, E. Perez, and our research greenhouse manager Linda Taylor for plant maintenance and assistance with this research in the greenhouse and in the lab. Several people made helpful suggestions on this manuscript including N. Diaz, M. Grasty, and J. Persinger. This research was supported by an NIH NIGMS BUILD EXITO grant (5TL4GM118965-03, 5UL1GM118964-03, and 5RL5GM118963-03) to Portland State University.

## Data Archiving

Data and all scripts for data analysis will be submitted to DataDryad upon publication.

## Appendix 1. Validity of the *δ*_*AD(k)*_ Estimate

For a pair of stems from the same genet, we assume that *w*_*G*_ represents the fitness of geitonogamous progeny after a cross between the pair of stems, and that *w*_*A*_ represents the fitness of autogamous progeny after selfing a single flower on one stem:

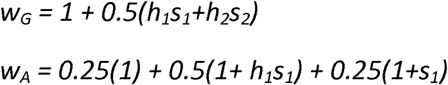

Where, *s*_1_ and *s*_2_ represent the selective effects of mutations, and *h*_1_ and *h*_2_ represent the dominance of single somatic mutations in stem 1 and stem 2, respectively.

For simulations, we randomly chose values of *s* ranging from -0.2 to 0.2 for 200 pairs of stems. Values of *h* for *s* > 0 were allowed to range from 0 to 0.7 for each stem. Because deleterious alleles tend to be recessive, values of h were constrained to range between 0 and 0.1 for *s* < 0. Dominance values were chosen assuming that deleterious somatic mutations with high levels of expression would be eliminated by cell lineage selection during vegetative growth, and beneficial mutations with high dominance would not display strong fitness effect differences between autogamous and geitonogamous progeny. To match our experimental design in *Mimulus guttatus*, we calculated the fitness of autogamous progeny (*w*_*A*_) for one of the stems in each pair. We then calculated *w*_*G*_ using information on s and h for both stems in each pair. Then we calculated the difference in fitness between each estimate as a measure of autogamy depression (*δ*_*AD(k)*_ = *w*_*A*_ – *w*_*G*_). This simulation was repeated 20 times (4,000 pairs of stems total) and average values were calculated.

### Results

1. Estimates of *δ*_*AD(k)*_ and *s*_1_ are strongly correlated, with R^2^ values ranging from 0.76 to 0.82 (one example of 200 random estimates):

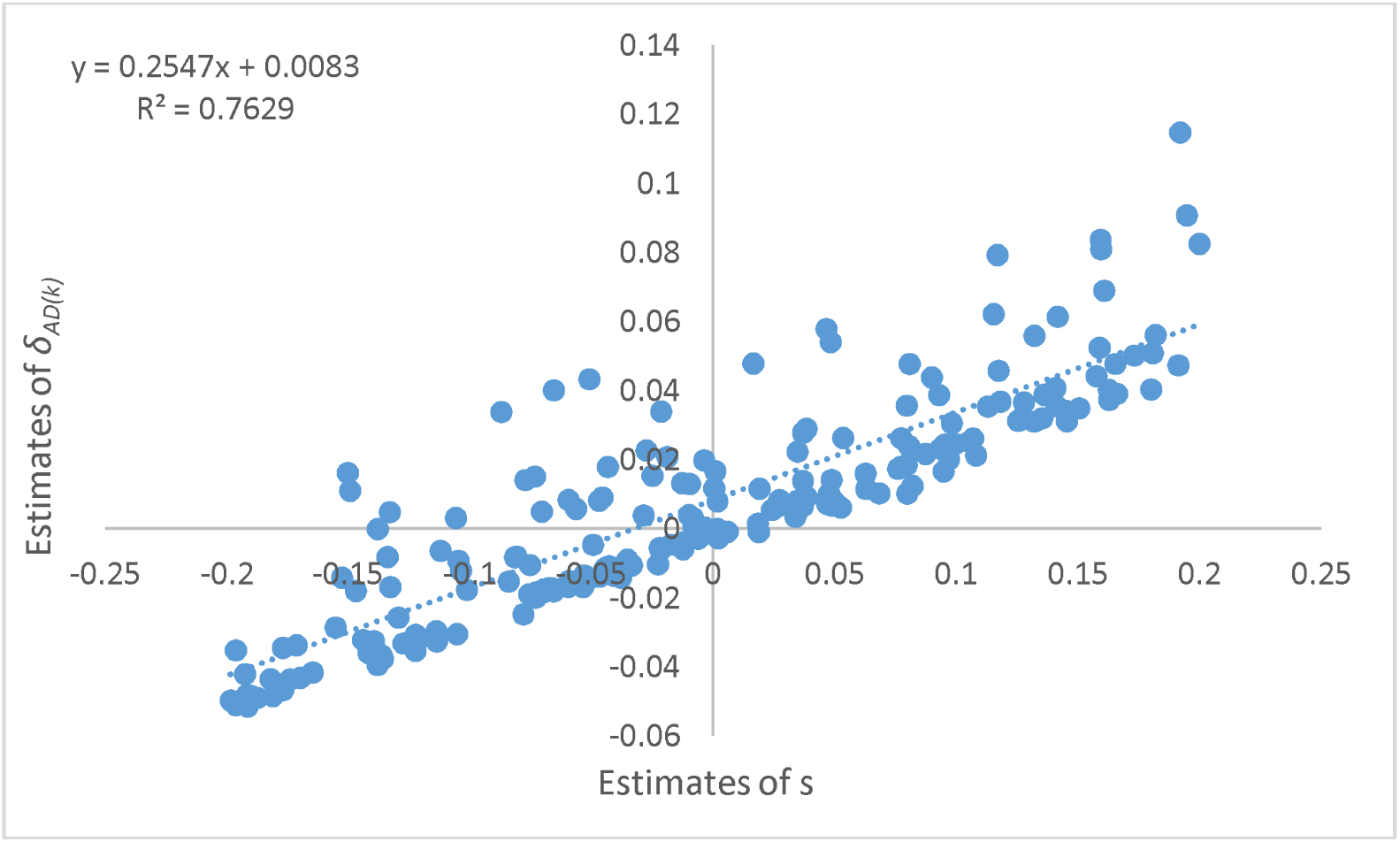
2. The frequency of *δ*_*AD(k)*_ estimates having the opposite sign of s_1_ was very low. Mean number of correct signs was 178.7 (+/- 1.04 standard error) and was 21.30 (+/-1.04; or 12%) for incorrect signs.
3. Estimates of *δ*_*AD(k)*_ with incorrect signs were closer to zero. Mean absolute values of s for *δ*_*AD(k)*_ estimates with incorrect signs was 0.054 (+/- 0.0021) and was 0.106 (+/-0.0011) for correct signs. Hence, estimates of *δ*_*AD(k)*_ that are further from zero are more reliable estimates of s_1_.

We note that the formulation used above for *w*_*G*_ could result in estimates that are out of range when large numbers of mutations are involved (i.e., *w*_*G*_ = *1 + 0*.*5 ∑h*_*i*_ *s*_*i*_ for a large number of loci segregating for deleterious alleles). A better estimate may be the average effect of *n* loci segregating for deleterious alleles (i.e. *∑h*_*i*_ *s*_*i*_ */n*). If we apply this approach for two loci (one mutation per stem) we obtain,

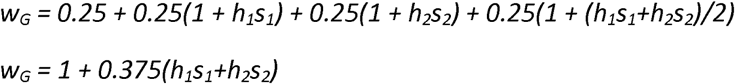

Using this formula, the relationship between δ_*AD(k)*_ and *s*_1_ is stronger (R^2^ ranging from 0.86 to 0.90), with only an average of 13 out of 200 estimates with opposite signs (7%). The average of correct signs was similar to the one above (0.105, +/- 0.0007), but the mean for incorrectly assigned values was much closer to zero (0.038, +/- 0.0022).

Another model for the effects of alleles at multiple loci on fitness assumes the interaction is multiplicative. In this case the estimate of *w*_*G*_ becomes,

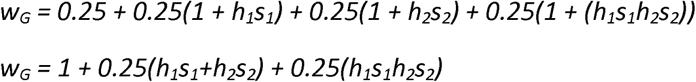

Using this formula, the relationship between δ_*AD(k)*_ and *s*_1_ is stronger (R^2^ ranging from 0.93 to 0.95), with only an average of 7.1 out of 200 estimates with opposite signs (4%). The average of correct signs was similar to the ones above (0.103, +/- 0.0010), but the mean for incorrectly assigned values was closer to zero (0.023, +/- 0.0020).

### Interpretation

These results indicate that our estimate of *δ*_*AD(k)*_ is a valid estimate of s for stem 1. However, it does appear that we are underestimating s (i.e., the slope of the line is less than 1), as these estimates do not consider the effects of multiple somatic mutations or possible genetic background effects (i.e. epistasis).

It is also notable that our simulation assumes a wider range of *h* values than are generally observed for deleterious mutations. Estimates indicate that the product of *hs* for deleterious alleles is generally around 0.02, and that the relationship between *s* and *h* is generally negative (hyperbolic; Lynch et al. 1999). Minor effects of deleterious mutations are expected for our data, because somatic mutations having strong effects in the heterozygous condition would be eliminated by cell lineage selection. Adding the constraint of a negative relationship between *h* and *s* to our simulation model would marginally improve the predictive power of *δ*_*AD(k)*_ for estimates of *s*.

These simulations also support our use of the correlation between w_A_ and w_G_ to confirm that our estimates are similar to the simulation estimates.

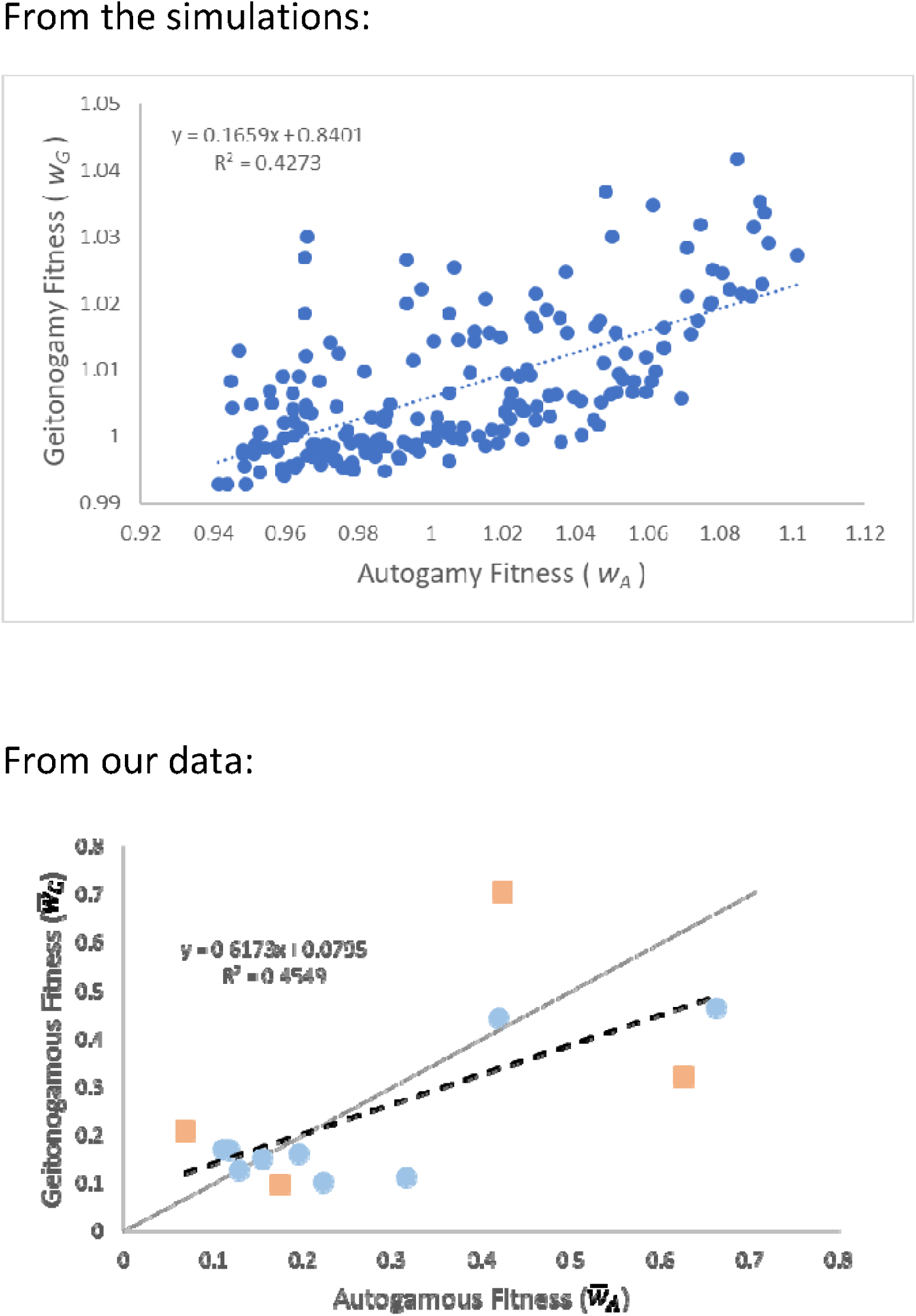

## Appendix 2. Estimating the selection coefficient due to somatic mutations from the variance in fitness among progeny

In this Appendix, we demonstrate that the standard deviation (SD) in fitness among progeny from autogamous crosses can be used to estimate the selection coefficient for the average effects of somatic mutations occurring within a stem. The variance in fitness among progeny from autogamy can be affected by the selection coefficient (*s*) and the dominance effects (*h*) of somatic mutations unique to a stem.

To evaluate the effects of a single mutation on progeny fitness, we assume fitnesses of;

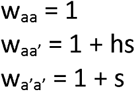

The mean progeny fitness is then calculated as

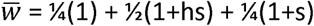

Which simplifies to

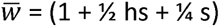

The variance in progeny fitness can be calculated as

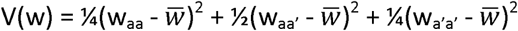

After substituting, we get

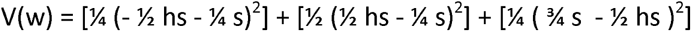

For simplification, we separate the effects of homozygous and heterozygous genotypes (terms within the three sets of square brackets)

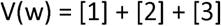

and evaluate each one;

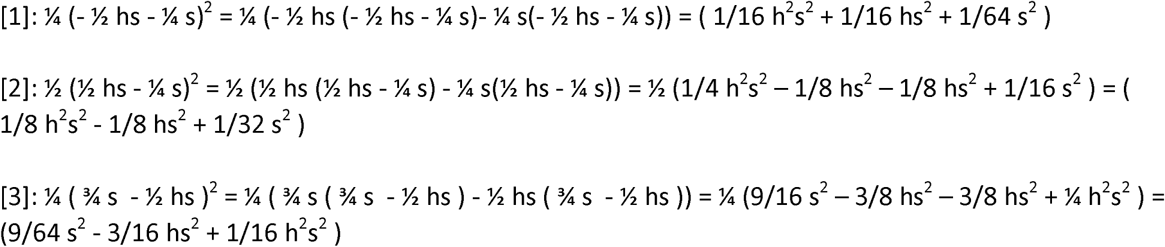

By combining and then simplifying these three results, we get

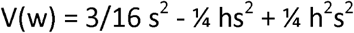

And the standard deviation can be obtained by taking the square root of V(w)

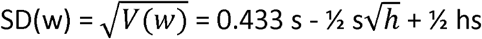

We note that this relationship holds regardless of the sign (positive or negative values of s), so for this relationship we can substitute *w* for *s*. The standard deviation in progeny fitness scales linearly with the selective effects of somatic mutations, and the effects of dominance on variation among progeny is minor (Fig. S1; which shows values of *s* or *w* predicted from values of SD for fitness ranging from 0 to 0.5). We note that the standard deviation is highest when mutations are completely recessive or dominant (h = 0 or 1) and SD(w) = 0.433s. When h = 0.5, the standard deviation is at its minimal value and SD(w) = 0.332s. By rearranging, we can obtain the regression equations to estimate the selective effects of somatic mutations from the standard deviation in progeny fitness as *s*_*SD*_ = *cSD*, where *c* is the slope of the regression line in Fig. S1. From this analysis, it is clear that the effects of *h* on the within-family SD in fitness is minor compared to the effects of *s*. We obtain estimates of the regression slopes from the standard deviation in progeny fitness to produce the relationships in Fig. S1;

**Fig. S1.**
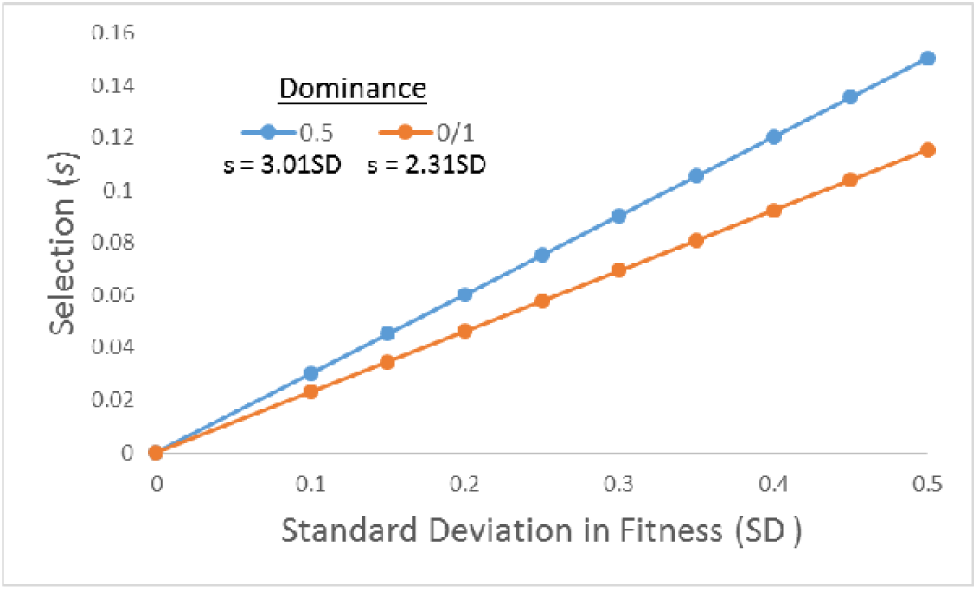
The effects of selection (*s*) or fitness (*w*) and dominance (*h*) on the standard deviation (SD) in fitness among progeny. The effect of dominance on variation in fitness is minimized when *h* = 0.5 and maximized when *h* = 0.0 and *h* = 1.0.

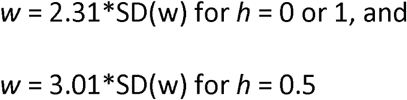

From these relationships we can estimate the selective effects of somatic mutations from the standard deviation in progeny fitness as *w*_*SD*_ = *cSD*, where *c* is the slope of the regression lines in Fig. S1.

## Appendix 3. Supplemental Tables and Figures

**Table S1.**

| Stem | <u>Autogamy</u> | | | <u>Geitonogamy</u> | | | $\delta_{AD(k)}$ | $\bar{w}_A - \bar{w}_G$ |
| --- | --- | --- | --- | --- | --- | --- | --- | --- |
| | $\bar{w}_A$ | N | Stdev | $\bar{w}_G$ | N | Stdev | | |
| JB51 | 0.407 | 33 | 0.171 | 0.144 | 7 | 0.080 | 0.263 | 0.078 |
| JB52 | 0.314 | 12 | 0.161 | 0.295 | 9 | 0.068 | 0.019 | -0.015 |
| JB54 | 0.258 | 8 | 0.066 | 0.298 | 11 | 0.103 | -0.040 | -0.071 |
| SMB11 | 0.286 | 22 | 0.072 | 0.260 | 9 | 0.077 | 0.009 | -0.043 |
| SMB51 | 0.256 | 11 | 0.054 | 0.323 | 4 | 0.035 | -0.067 | -0.073 |
| SMB52 | 0.273 | 7 | 0.072 | 0.280 | 6 | 0.051 | -0.007 | -0.056 |
| SMB53 | 0.245 | 4 | 0.021 | 0.327 | 6 | 0.108 | -0.081 | -0.083 |
| SMB54 | 0.289 | 10 | 0.023 | 0.245 | 5 | 0.056 | 0.044 | -0.040 |
| SMC11 | 0.303 | 16 | 0.071 | 0.238 | 8 | 0.085 | 0.065 | -0.026 |
| SMC12 | 0.356 | 8 | 0.071 | 0.239 | 6 | 0.059 | 0.117 | 0.027 |
| C12 | 0.423 | 36 | 0.170 | 0.707 | 5 | 0.153 | -0.283 | 0.094 |
| C13 | 0.626 | 9 | 0.195 | 0.322 | 7 | 0.237 | 0.090 | 0.297 |
| D17 | 0.692 | 5 | 0.170 | 0.485 | 4 | 0.314 | 0.207 | 0.363 |
| D18 | 0.419 | 12 | 0.130 | 0.443 | 13 | 0.211 | -0.025 | 0.090 |

**Table S2.** Analysis of variance results for seed set after autogamous and geitonogamous pollinations (Cross) with limited and excess pollen (Pollen). Population (Pop), maternal stem nested within population (Mat(Pop)), and pollination date (Date) were declared as random effects in the model.

| Source | DF | Sum of Squares | Mean Square | F Value | Pr > F |
| --- | --- | --- | --- | --- | --- |
| **Model** | 33 | 21623.26022 | 655.25031 | 1.43 | 0.0888 |
| **Cross** | 1 | 100.647606 | 100.647606 | 0.22 | 0.6400 |
| **Pop** | 2 | 552.228601 | 276.114301 | 0.60 | 0.5487 |
| **Pollen** | 1 | 2183.436724 | 2183.436724 | 4.78 | 0.0312 |
| **Mat(Pop)** | 18 | 6490.364191 | 360.575788 | 0.79 | 0.7089 |
| **Date** | 11 | 8000.908005 | 727.355273 | 1.59 | 0.1127 |
| **Error** | 101 | 46182.07311 | 457.24825 |  |  |

**Table S3.** Analysis of variance results for aborted ovules after autogamous and geitonogamous pollinations (Cross) with limited and excess pollen (Pollen). Population (Pop), maternal stem nested within population (Mat(Pop)), and pollination date (Date) were declared as random effects in the model.

| Source | DF | Sum of Squares | Mean Square | F Value | Pr > F |
| --- | --- | --- | --- | --- | --- |
| **Model** | 33 | 9337.61468 | 282.95802 | 2.76 | <.0001 |
| **Cross** | 1 | 449.176948 | 449.176948 | 4.38 | 0.0389 |
| **Pop** | 2 | 2174.689160 | 1087.344580 | 10.60 | <.0001 |
| **Pollen** | 1 | 0.597410 | 0.597410 | 0.01 | 0.9393 |
| **Mat(Pop)** | 18 | 2217.350757 | 123.186153 | 1.20 | 0.2751 |
| **Date** | 11 | 1995.681210 | 181.425565 | 1.77 | 0.0695 |
| **Error** | 101 | 10363.24458 | 102.60638 |  |  |

**Table S4.** Analysis of variance for progeny fitness (biomass weighted by survival of seedlings) after autogamous and geitonogamous pollinations (Cross) for ten maternal stems (Mat) and their interaction (Mat*Cross). Planting block (Tray) was declared as a random effect in the model.

| Source | DF | Sum of Squares | Mean Square | F Value | Pr > F |
| --- | --- | --- | --- | --- | --- |
| **Model** | 33 | 1.42418594 | 0.04315715 | 4.18 | <.0001 |
| **Tray** | 14 | 0.45040769 | 0.03217198 | 3.11 | 0.0002 |
| **Mat** | 9 | 0.06845581 | 0.00760620 | 0.74 | 0.6756 |
| **Cross** | 1 | 0.03196532 | 0.03196532 | 3.09 | 0.0805 |
| **Mat\*Cross** | 9 | 0.51373985 | 0.05708221 | 5.52 | <.0001 |
| **Error** | 168 | 1.73638043 | 0.01033560 |  |  |

**Table S5.** Analysis of variance for progeny fitness (biomass weighted by survival of seedlings) after autogamous and geitonogamous pollinations (Cross) for seven paired pollinations at individual nodes (Stem/Node) and their interaction (Stem/Node*Cross). Hydroponic planting block (Tray) was declared as a random effect in the model and initial size of seedlings (Initial Size) was entered as a covariate.

| Source | DF | Sum of Squares | Mean Square | F Value | Pr > F |
| --- | --- | --- | --- | --- | --- |
| **Model** | 23 | 1.31384069 | 0.05712351 | 6.05 | <.0001 |
| **Tray** | 11 | 0.46856600 | 0.04259691 | 4.51 | <.0001 |
| **Initial Size** | 1 | 0.46861414 | 0.46861414 | 49.61 | <.0001 |
| **Stem/Node** | 6 | 0.12071536 | 0.02011923 | 2.13 | 0.0514 |
| **Cross** | 1 | 0.00555294 | 0.00555294 | 0.59 | 0.4441 |
| **Stem/Node\*Cross** | 4 | 0.13005812 | 0.03251453 | 3.44 | 0.0095 |
| **Error** | 203 | 1.91760656 | 0.00944634 |  |  |

**Table S6.**
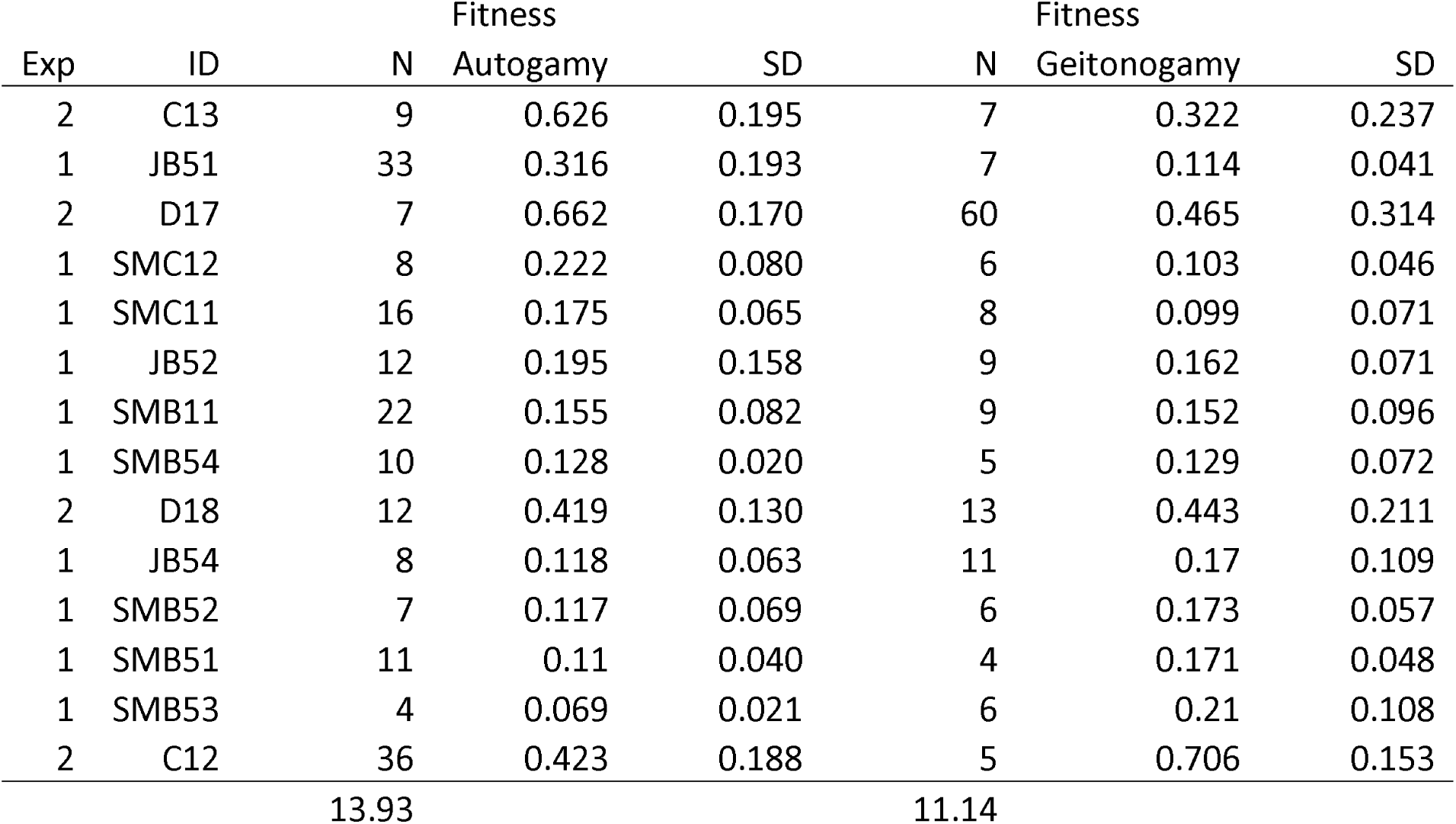
Means and standard deviations (SD) for progeny groups from autogamous and geitonogamous pollinations in experiments 1 and 2. Experiment 1 included ten different genets. Experiment 2 was conducted with two ramets (C1 and D1) from the same genet. The fitness of progeny was estimated as its final biomass (log transformed to improve normality), weighted by the survival frequency of progeny from the same cross.

**Fig. S2.**
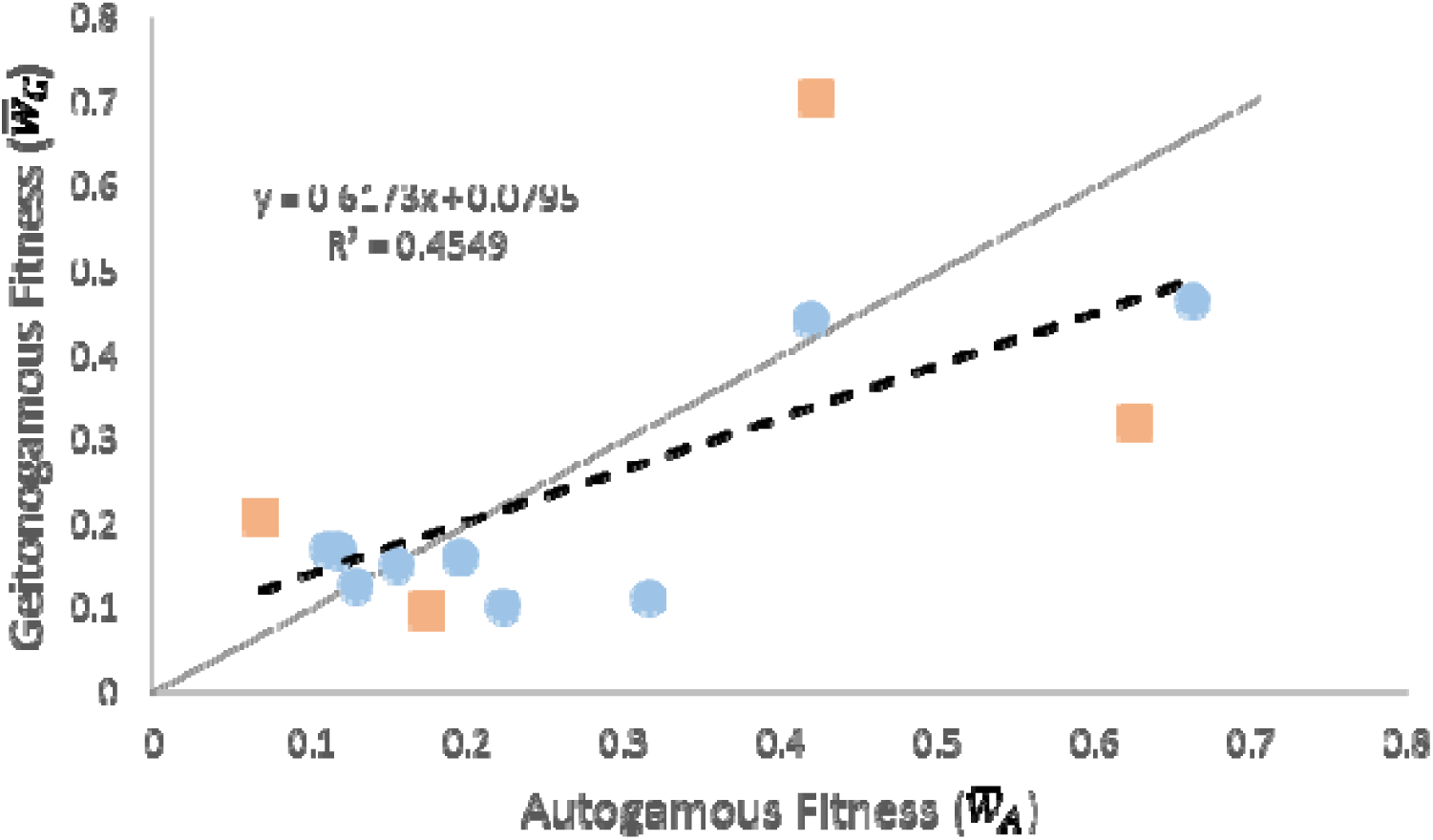
The relationship between the autogamous and geitonogamous progeny from the 14 comparisons from experiments 1 (blue circles) and 2 (orange squares) as listed in Table S6. The solid line represents a 1:1 relationship and the dashed line represents the best-fit to the data. Stems falling above the solid line apparently had a prevalence of deleterious somatic mutations whereas those falling below the line displayed a prevalence of beneficial somatic mutations.

**Fig. S3.**
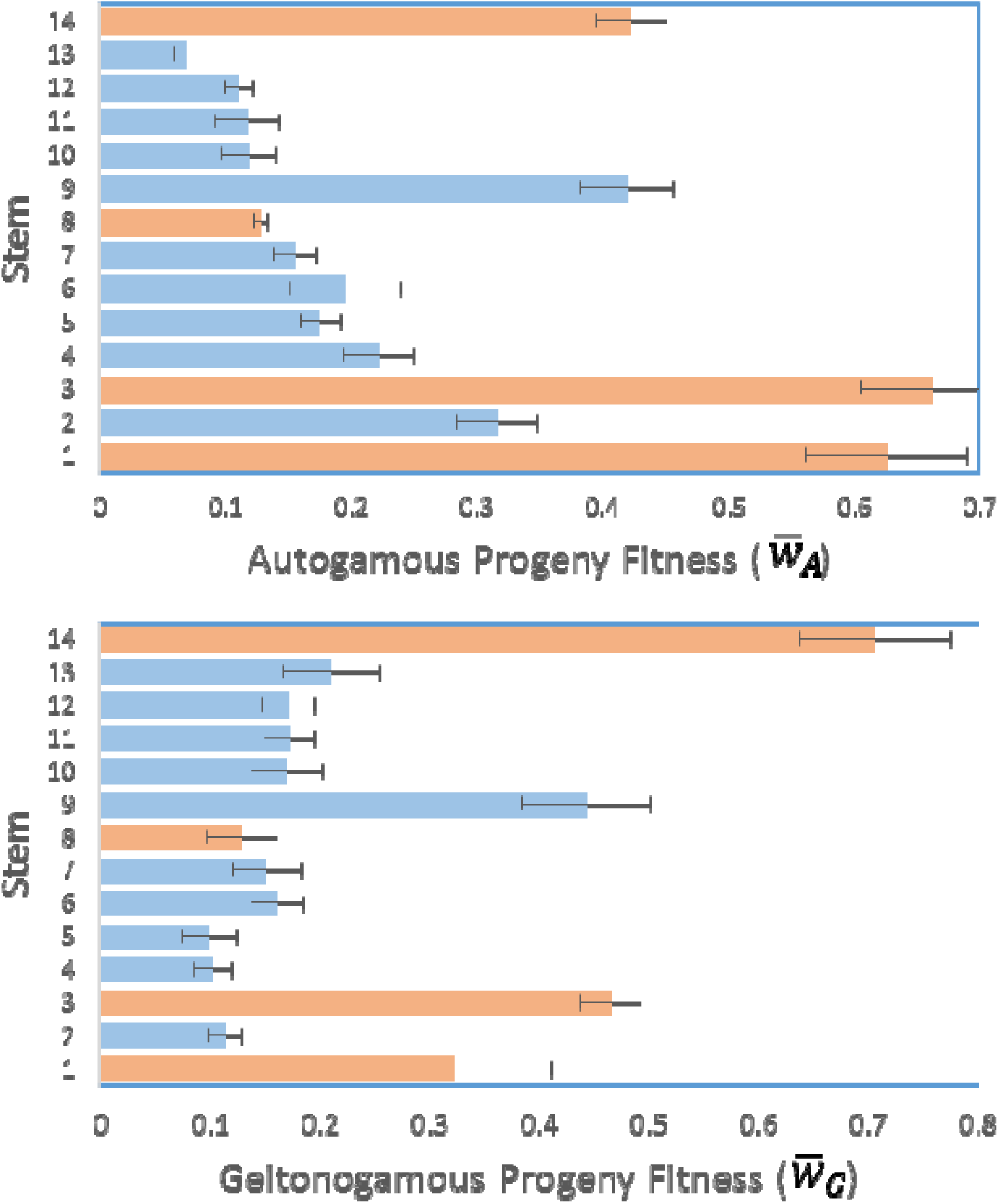
Means and standard errors (lines) for fitness estimates of autogamous (upper panel) and geitonogamous (lower panel) progeny groups from experiment 1 (blue bars) and 2 (orange bars) listed in Table S6. Progeny groups are ordered as in Fig. 4.

